# Temperature drives Zika virus transmission: evidence from empirical and mathematical models

**DOI:** 10.1101/259531

**Authors:** Blanka Tesla, Leah R. Demakovsky, Erin A. Mordecai, Sadie J. Ryan, Matthew H. Bonds, Calistus N. Ngonghala, Melinda A. Brindley, Courtney C. Murdock

**Author notes:** Corresponding author **(CM)**.

## Abstract

Temperature is a strong driver of vector-borne disease transmission. Yet, for emerging arboviruses we lack fundamental knowledge on the relationship between transmission and temperature. Current models rely on the untested assumption that Zika virus responds similarly to dengue virus, potentially limiting our ability to accurately predict the spread of Zika. We conducted experiments to estimate the thermal performance of Zika virus (ZIKV) in field-derived *Aedes aegypti* across eight constant temperatures. We observed strong, unimodal effects of temperature on vector competence, extrinsic incubation period, and mosquito survival. We used thermal responses of these traits to update an existing temperature-dependent model to infer temperature effects on ZIKV transmission. ZIKV transmission was optimized at 29°C, and had a thermal range of 22.7°C - 34.7°C. Thus, as temperatures move toward the predicted thermal optimum (29°C) due to climate change, urbanization, or seasonally, Zika could expand north and into longer seasons. In contrast, areas that are near the thermal optimum were predicted to experience a decrease in overall environmental suitability. We also demonstrate that the predicted thermal minimum for Zika transmission is 5°C warmer than that of dengue, and current global estimates on the environmental suitability for Zika are greatly over-predicting its possible range.

## Introduction

Mosquito-borne viruses are an emerging threat impacting human health and well-being. Epidemics of dengue (DENV), chikungunya, and Zika (ZIKV) have spilled out of Africa to spread explosively throughout the world creating public health crises. Worldwide, an estimated 3.9 billion people living within 120 countries are at risk [1]. In 2015-2016, ZIKV spread throughout the Americas including the continental U.S., resulting in over 360,000 suspected cases, with likely many more undetected [2]. With the rise of neurological disorders and birth defects, such as Guillain-Barré and congenital Zika virus syndrome [3, 4], ZIKV became widely feared and was declared a “public health emergency of international concern” by the World Health Organization in 2016 [5]. In spite of growing research efforts to develop new therapeutics, vaccines, and innovative mosquito control technologies, mitigating arbovirus disease spread still depends on conventional mosquito control methods and public education. Thus, substantial efforts have been made to predict who ZIKV will spread seasonally, geographically, and with the effects of climate change [e.g. 6, 7–9].

There are several key gaps that potentially affect our ability to predict, and ultimately, mitigate the factors that influence transmission risk and arbovirus emergence globally. First, current models predicting mosquito distributions or virus transmission are often limited by a relatively poor understanding of the relationships among mosquito vectors, pathogens, and the environment. There is substantial evidence that temperature variability is a key driver of disease transmission across diverse vector-borne pathogen systems [8, 10–15]. Mosquitoes are small ectothermic animals and their physiology [16–18], life history [8, 19], and vectorial capacity [10, 20–22] exhibit unimodal responses to changes in temperature. Transmission depends in large part on the ability of mosquitoes to survive the extrinsic incubation period (EIP), become infectious, and bite new hosts, so differential (unimodal) impacts of temperature on survival, vector competence, and EIP have highly nonlinear effects on transmission. Warmer temperatures do not necessarily translate into more infectious mosquitoes [8, 20, 23]. Second, current models often ignore the low quality and quantity of existing data. Even in systems that are fairly well-studied (e.g. *Plasmodium falciparum* and DENV), key parameters are often estimated from only a few studies. Finally, current transmission models often assume, with little justification, that the relationship between temperature and EIP is monotonic [24], or that the relationships between temperature, EIP, and vector competence of less-studied arboviruses (e.g. chikungunya and ZIKV) are similar to DENV [8, 9, 25].

To advance our fundamental scientific understanding of the relationship between temperature and ZIKV transmission, we conducted a series of experiments to estimate the thermal performance of ZIKV (vector competence, the extrinsic incubation rate, and the daily per capita mosquito mortality rate) in field-derived *Ae. aegypti* across eight different constant temperatures ranging from 16 - 38°C. We fit a series of nonlinear functions to estimate the thermal responses of the above traits. These thermal responses were incorporated into a temperature dependent basic reproductive number (*R_0_*) model developed for *Ae. aegypti* and DENV [14] to infer how temperature variation will impact ZIKV transmission.

## Methods

### Experimental mosquito infections and forced salivations

For details on virus culture and mosquito rearing, see supplementary information (Text

S1). For each biological replicate, we separated 8,000 1 to 3-day-old females (field derived *Ae. aegypti*, F4 generation) prior to ZIKV infection (Fig S1). Mosquitoes were kept in 64 oz. paper cups and provided with water, which was withdrawn 12 hours before feeding. We offered them either an infectious blood meal containing ZIKV at a final concentration of 10^6^ PFU/mL or an uninfected, control blood meal. The blood meal was prepared by washing human blood three times in RPMI medium and the pelleted red blood cells (50%) were resuspended in 33% DMEM, 20% FBS, 1% sucrose, and 5 mmol/L ATP. For the infectious blood meal, we mixed the blood mixture with ZIKV diluted in DMEM (2*10^6^ PFU/mL) at a 1:1 ratio. Mosquitoes were blood-fed through a water-jacketed membrane feeder for 30 min, after which we randomly distributed 2,000 ZIKV-exposed engorged mosquitoes and 2,000 unexposed blood-fed control mosquitoes into mesh-covered paper cups (250 mosquitoes per cup). We then placed one ZIKV-exposed and one control cup at each temperature treatment (Percival Scientific): 16°C, 20°C, 24°C, 28°C, 32°C, 34°C, 36°C, and 38°C ± 0.5°C. Chambers were set to 80% + 5% relative humidity and a 12:12 light:dark cycle, and mosquitoes were maintained on 10% sucrose for the duration of the experiment. Mosquito mortality was monitored and recorded daily.

Every three days (up to day 21) we force-salivated 20 ZIKV-exposed mosquitoes per treatment group by immobilizing mosquitoes on ice, removing their legs and wings, and placing the proboscis of each mosquito into a pipet tip (containing 35 μL FBS with 3 mmol/L ATP) for 30 min on a 35°C warming plate. After salivation, we collected mosquito saliva, heads and legs, and bodies into 700 μL of DMEM with 1x antibiotic/antimycotic. Each tissue was homogenized in a QIAGEN TissueLyzer at 30 cycles/second for 30 seconds, and centrifuged at 17,000xg for 5 minutes at 4°C. To measure the proportion of mosquitoes that became infected, disseminated infection, and became infectious at each temperature, we tested for the presence/absence of ZIKV in mosquito bodies, legs and heads, and saliva, respectively, using plaque assays on Vero cells as described above. Two full biological replicates were performed (Fig S1).

### Statistical analysis

The effects of temperature was assessed on four different metrics of ZIKV infection. We used the numbers of mosquitoes becoming infected (ZIKV positive bodies), disseminated (ZIKV positive legs / heads), and infectious (ZIKV positive saliva) out of total numbers of mosquitoes exposed to assess the effect of temperature on the likelihood of ZIKV infection, dissemination, and infectiousness at the population level. We also used the numbers of mosquitoes that became infectious out of those successfully infected (positive bodies) as a measure of ZIKV dissemination efficiency. For each response variable, we used generalized linear mixed models (IBM^®^ SPSS^®^ Statistics 1.0.0.407), normal distribution and identity link function, to estimate the effects of temperature (16°C, 20°C, 24°C, 28°C, 32°C, 34°C, 36°C, 38°C), days post infection (dpi 3, 6, 9, 12, 15, 18, 21), and the interaction between temperature and dpi (fixed factors). Mosquito batch nested within experimental replicate was included in all models as a random factor. We determined the best model fit and distributions based on Akaike Information Criterion (AIC), the dispersion parameter, and by plotting model residuals. Sequential Bonferroni tests were used to assess the significance of pairwise comparisons within a significant main effect or interaction, and *p*-values greater than 0.05 were considered non-significant. Finally, to estimate the effects of temperature, ZIKV exposure and the interaction between temperature and ZIKV exposure on the daily probability of mosquito survival, we used the same framework in a Cox proportional hazards model (SAS^®^ Studio, 3.6 Basic Edition) with temperature, infection status (ZIKV-exposed or control) and the interaction as fixed factors, with mosquito batch nested within experimental replicate as a random factor.

### Mechanistic *R_0_* model

In previous work, we assembled trait thermal response estimates from laboratory experiments that manipulated temperature and measured each of the following traits for *Ae. aegypti* and DENV: egg-to-adult development rate (*MDR*), survival probability (*p_EA_*), fecundity (*EFD*; eggs per female per day), biting rate (*a*), adult mosquito mortality rate (*μ*), extrinsic incubation rate (*EIR*), and vector competence (*bc*; equal to the proportion of exposed mosquitoes that become infected times the proportion of infected mosquitoes that become infectious, with virus in their saliva). We then synthesized them into an estimate for the thermal response of *R_0_*, the expected number of new cases generated by a single infectious person or mosquito introduced into a fully susceptible population throughout the period within which the person or mosquito is infectious [8]:

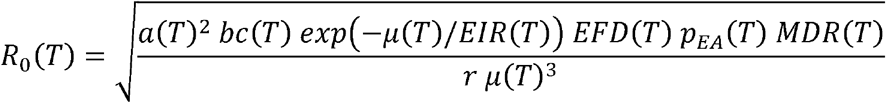

where *r* is the human recovery rate, *T* is environmental temperature, and *T* attached to a parameter indicates that the parameter is dependent on temperature. Here, we update three of these thermal response functions—average adult mosquito lifespan (*lf=1/μ*), extrinsic incubation rate (*EIR*), and vector competence (*bc*) —using the new experimental data from *Ae. aegypti* mosquitoes exposed to ZIKV-infected blood meals across a range of constant temperatures (see Text S1).

### Mapping Seasonal Transmission Range

To illustrate predicted temperature suitability for Zika transmission in the Americas, we mapped the number of months for which *R_0_*(*T*)>0 for the posterior median response, based on the temperature-dependent model derived here and previously [8]. We calculated *R_0_*(*T*) at 0.1°C increments and projected it onto the landscape for monthly mean current temperatures from WorldClim data at a 5-minute resolution (approximately 10km^2^ at the equator). Climate data layers were extracted for the geographic area and defined using the Global Administrative Boundaries Databases [26]. All map calculations and manipulations were run in R using packages ‘raster’ [27], ‘maptools’ [28], and ‘Rgdal’ [29]. Resulting GeoTiffs were rendered in ArcGIS 10.3.1 [30], and mapped as figures. We then used the area represented by 6 months and 12 months of transmission suitability to calculate and display the difference between a previous model parameterized on the *Ae. aegypti-DENV* system [8] and our current predictions.

## Results

We found significant effects of temperature, days post-infection (dpi), and the interaction on the number of mosquitoes that became infected (ZIKV-positive bodies), that disseminated infection (ZIKV-positive legs and heads), and that became infectious (ZIKV-positive saliva). We also found significant effects of temperature, dpi, and the interaction on the overall transmission efficiency of ZIKV. Finally, these effects translated into significant effects of temperature on *R_0_*, or predicted risk of transmission for ZIKV, which differed from previous estimates generated from DENV specific models.

### The effect of temperature on ZIKV infection and infection dynamics

We observed strong, unimodal effects of temperature on the number of mosquitoes infected, with disseminated infections, and that became infectious (Table 1, Fig 1). While all three response variables dropped at both cool and warm temperatures, this decrease was more pronounced as the infection progressed (Fig 1). For example, the likelihood of becoming infected was the most permissive to temperature variation, with the number of infected mosquitoes minimized at 16°C (6%), maximized from 24°C-34°C (75% - 89%), and again minimized at 38°C (7%). The likelihood of viral dissemination was more constrained, with numbers of mosquitoes with disseminated infections minimized at 16-20°C (4% - 3%), maximized at 28-34°C (65% - 77%), and again minimized at 38°C (5%). Finally, the likelihood of mosquitoes becoming infectious was the most sensitive to temperature, with the numbers of infectious mosquitoes minimized from 16-24°C (0%-4%), maximized between 28-34°C (23%-19%), and again minimized from 36-38°C (5%-0.4%). We also observed a significant effect of temperature on the rate that virus disseminated through the mosquito and could be detected in saliva (*temperature x day*, Table 1, Fig 2). In general (with the exception of 36°C and 38°C), we observed increases in the numbers of mosquitoes with ZIKV positive bodies, legs and heads, and saliva with time (Fig 2) suggesting that the time at which ZIKV was detected in these samples decreased with increasing temperature. At 36°C and 38°C, we see declines in these response variables over time due to high mosquito mortality.

**Fig 1.**
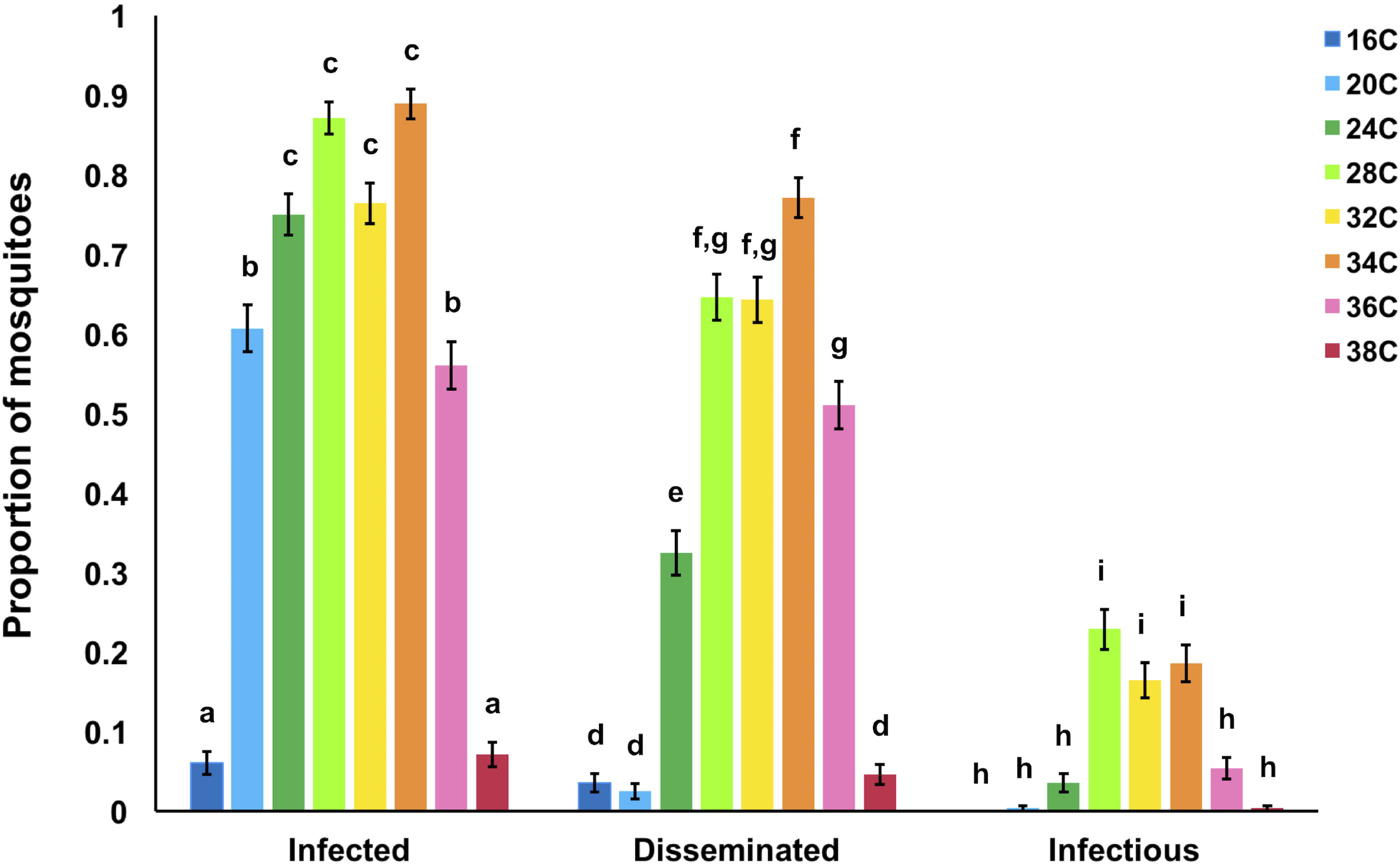
Temperature effect on the proportion of mosquitoes infected, with disseminated infections, and infectious. The effect of eight different constant temperatures (16°C, 20°C, 24°C, 28°C, 32°C, 34°C, 36°C, 38°C) on the proportion of mosquitoes infected (ZIKV positive bodies compared to total number of exposed), with disseminated infections (ZIKV positive heads compared to total number exposed), and infectious (ZIKV positive saliva compared to total number exposed). Results with no common letters were significantly different (*p* ≤ 0.05).

**Fig 2.**
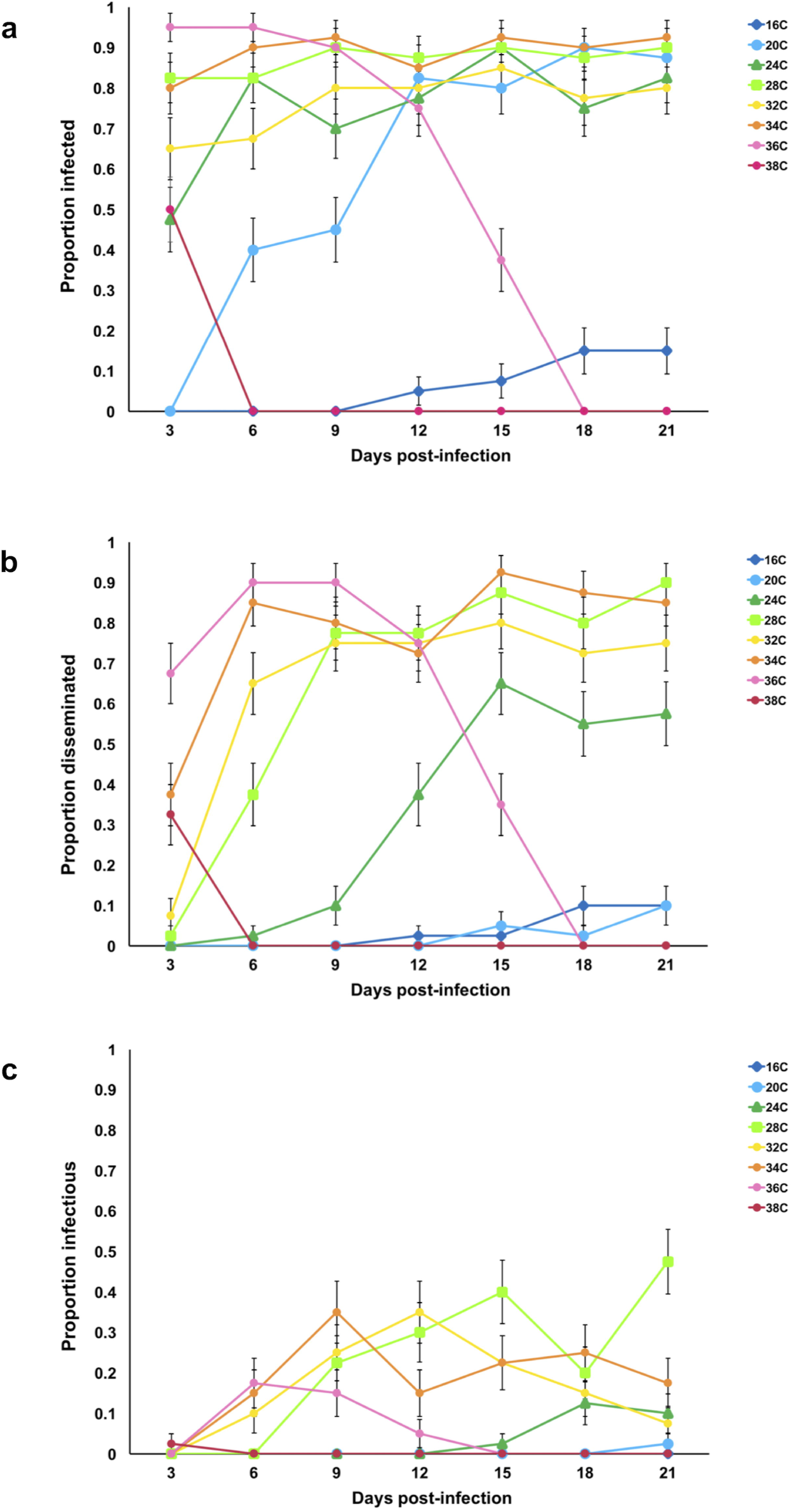
Days post-infection and the proportion of mosquitoes infected, with disseminated infections, and infectious. The relationship between days post-infection (3, 6, 9, 12, 15, 18, 21) and the proportion of mosquitoes infected (A, ZIKV positive bodies), with disseminated infections (B, ZIKV positive legs and heads), and infectious (C, ZIKV positive saliva) out of the total mosquitoes exposed to ZIKV at eight different constant temperatures (16°C, 20°C, 24°C, 28°C, 32°C, 34°C, 36°C, 38°C).

**Table 1.** Results from generalized linear mixed effects models examining the effects of temperature, day, and the interaction on the numbers (out of total exposed) of mosquitoes infected, with disseminated infections, infectiousness, and a measure of dissemination efficiency.

| response variables | temperature |  |  | day |  |  | temperature x day |  |  |
| --- | --- | --- | --- | --- | --- | --- | --- | --- | --- |
|  | F | d.f. | p-value | F | d.f. | p-value | F | d.f. | p-value |
| number infected | 86.159 | 7 | **<0.0001** | 1.349 | 6 | 0.251 | 8.374 | 42 | **<0.0001** |
| number disseminated | 139.085 | 7 | **<0.0001** | 14.742 | 6 | **<0.0001** | 12.477 | 42 | **<0.0001** |
| number infectious | 34.012 | 7 | **<0.0001** | 8.876 | 6 | **<0.0001** | 4.846 | 42 | **<0.0001** |
| dissemination efficiency | 15.308 | 7 | **<0.0001** | 7.699 | 6 | **<0.0001** | 4.431 | 42 | **<0.0001** |

**Table 2.** The time (days post infection, dpi) required for first detection and to reach the plateau of numbers of mosquitoes infected, with disseminated infections, and infectious.

| temperature | 16°C | 20°C | 24°C | 28°C | 32°C | 34°C | 36°C | 38°C |
| --- | --- | --- | --- | --- | --- | --- | --- | --- |
| time (dpi) to first detection of: |  |  |  |  |  |  |  |  |
| infection | 12 | 6 | 3 | 3 | 3 | 3 | 3 | 3 |
| dissemination | 12 | 15 | 6 | 3 | 3 | 3 | 3 | 3 |
| infectiousness |  | 21 | 15 | 9 | 6 | 6 | 6 | 3 |
| time (dpi) to reach plateau of: |  |  |  |  |  |  |  |  |
| infection | 18 | 12 | 6 | 3 | 9 | 3 | 3 | 3 |
| dissemination | 18 | 21 | 15 | 15 | 9 | 6 | 6 | 3 |
| infectiousness |  | 21 | 18 | 15 | 12 | 9 | 6 | 3 |

### The effects of temperature on ZIKV dissemination efficiency

We observed significant effects of temperature, dpi, and the interaction on the overall dissemination efficiency of ZIKV. ZIKV dissemination efficiency was maximized from 28 – 34°C suggesting that ZIKV escape from the midgut and salivary gland invasion was most efficient at these temperatures (Fig 3). In contrast, dissemination efficiency was minimized at both cooler (16 - 20°C) and warmer temperatures (38°C). Interestingly, cooler temperatures had a more dramatic effect on dissemination efficiency relative to warmer temperatures. For example, although 60% of exposed mosquitoes became successfully infected at 20°C, we had very low salivary gland invasion, with only one mosquito across both trials becoming infectious. In contrast, at warm temperatures infection and dissemination efficiencies were very robust (Fig S2), but the mortality associated with the warm temperatures resulted in low numbers of mosquitoes that were capable of being infectious. Finally, of those successfully infected, we observed successful salivary gland invasion to occur earlier in the infection process as temperatures warmed (Fig 3).

**Fig 3.**
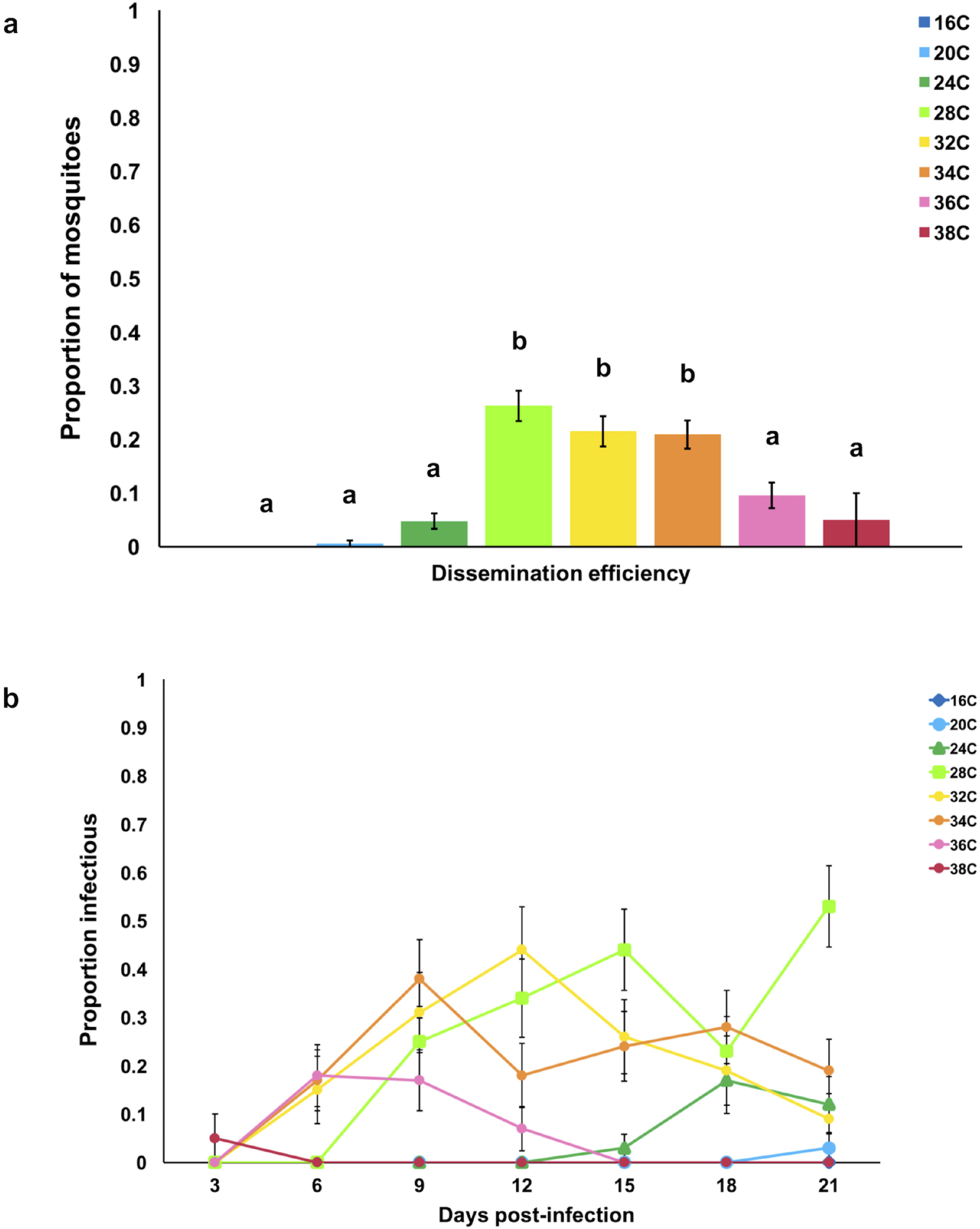
Temperature effect on the dissemination efficiency. (A) The effect of eight different constant temperatures (16°C, 20°C, 24°C, 28°C, 32°C, 34°C, 36°C, 38°C) and (B) days post-infection (3, 6, 9, 12, 15, 18, 21) on the dissemination efficiency (proportion of ZIKV positive saliva relative to positive bodies). Results with no common letters were significantly different (*p* ≤ 0.05).

### The effect of temperature on mosquito survival

We observed significant effects of temperature and an interaction between temperature and ZIKV exposure on the daily probability of mosquito survival (Fig S3, Table 3). Overall, the daily probability of mosquito survival was highest for mosquitoes housed at 24°C and 28°C relative to cooler (16 - 20°C) and warmer (32 - 38°C) temperatures. Mosquito survival was lowest at the warmest temperature of 38°C, with no mosquitoes surviving past 3 dpi. ZIKV-exposed mosquitoes experienced a higher daily probability of survival at 24°C and 28°C relative to unexposed, control mosquitoes with greater than 90% survival at the optimal temperatures.

**Table 3.**
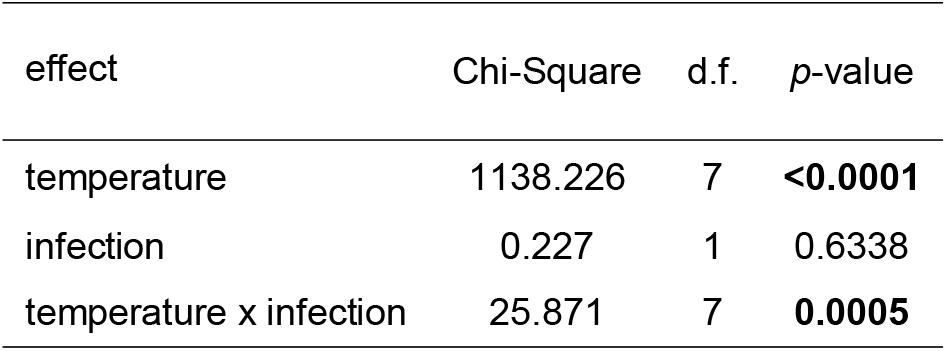
Results from Cox mixed-effects model examining the effects of temperature (16°C, 20°C, 24°C, 28°C, 32°C, 34°C, 36°C, 38°C), infection status (exposed or not exposed) and the interaction on the daily probability of mosquito survival.

| effect | Chi-Square | d.f. | p-value |
| --- | --- | --- | --- |
| temperature | 1138.226 | 7 | **<0.0001** |
| infection | 0.227 | 1 | 0.6338 |
| temperature x infection | 25.871 | 7 | **0.0005** |

### The effect of temperature on ZIKV transmission risk

Trait thermal responses for lifespan, vector competence, and extrinsic incubation rate were all unimodal (Fig 4, Table 1 SI, Fig S4). Mosquito lifespan and vector competence thermal responses were symmetrical, peaking at 24.2°C (95% CI: 21.9 – 25.9°C) and 30.6°C (95% CI: 29.6 – 31.4°C), respectively, while the extrinsic incubation rate thermal response was asymmetrical with a peak at 36.4°C (95% CI: 33.6 – 39.1°C). Applying these new trait thermal responses to the *R_0_*(*T*) model [8], we found that *R_0_*(*T*) peaked at 28.9°C (95% CI: 28.1 – 29.5°C), with lower and upper limits of 22.7°C (95% CI: 21.0 – 23.9°C) and 34.7°C (95% CI: 34.1 – 35.8°C), respectively (Fig 5). The seasonal transmission of ZIKV was predicted to be more constricted in latitudinal range from this temperature –transmission relationship than what has been predicted previously [8], primarily because the predicted thermal minimum for ZIKV was 5°C warmer than for DENV (S5 Fig). The estimated change in land area this represents in the Americas for endemic (12 month, year-round suitability), and overall predicted range (1-12 months suitability) is 4.3 million km^2^ and 6.03 million km^2^, respectively (Fig 6).

**Fig 4.**
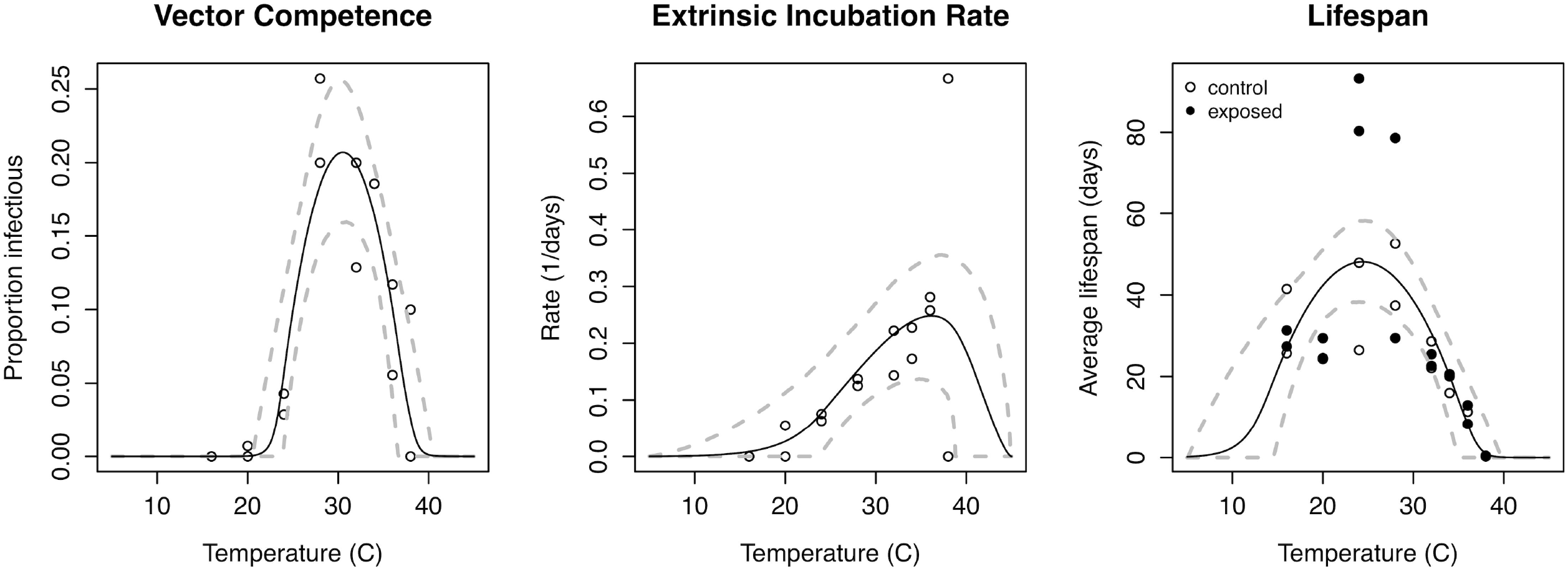
Effect of temperature and estimated vector competence, extrinsic incubation rate and mosquito lifespan. Trait thermal responses, fit from laboratory experimental data. Vector competence (left), is the maximum proportion of virus-exposed mosquitoes with virus in their saliva, across temperatures and replicates. Extrinsic incubation rate (middle) is the inverse of the time required to reach half of the maximum proportion infectious (days^−1^) for each temperature and replicate. Lifespan is the average lifespan of mosquitoes in each temperature and replicate (days), shown in filled (virus-exposed) and open (sham-inoculated) points. Solid lines represent posterior means; dashed lines represent 95% credible intervals.

**Fig 5.**
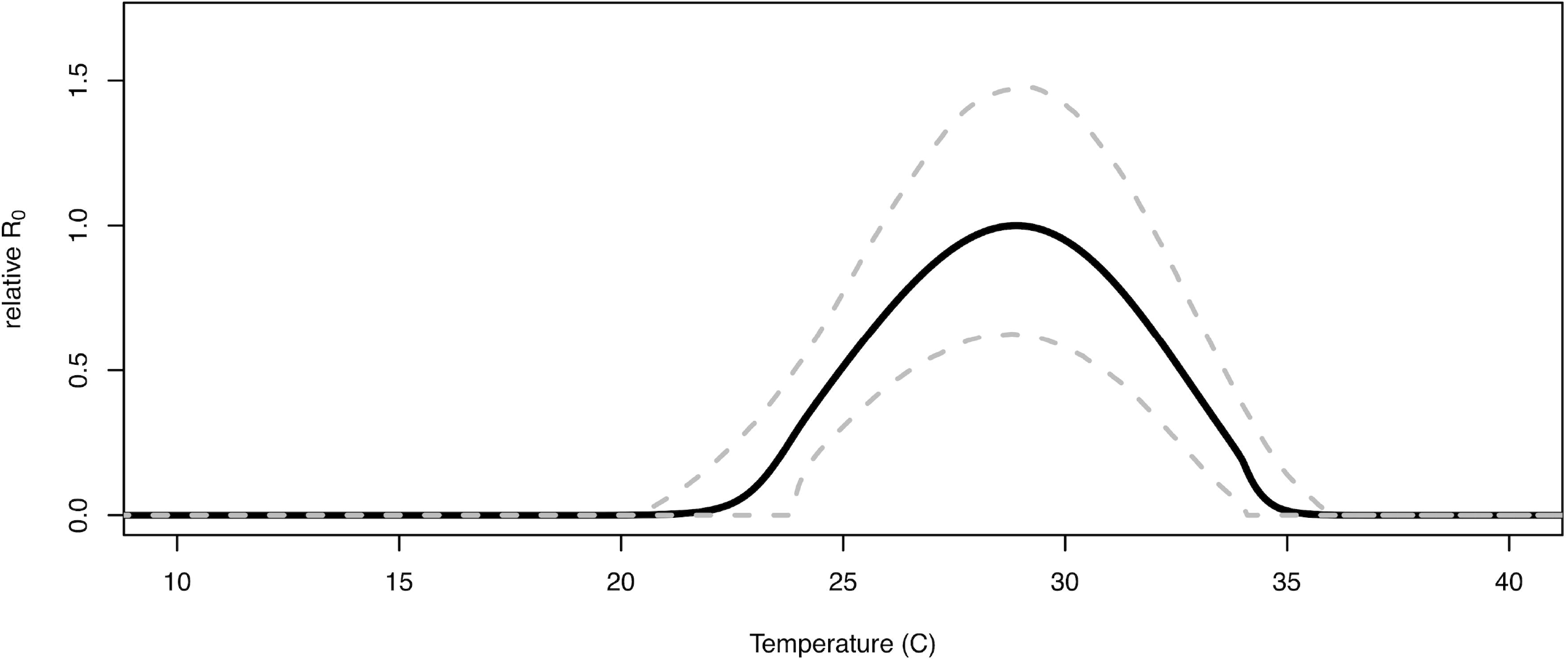
Effect of temperature on *R_0_*. Effect of temperature on *R_0_* (top). Solid line is the mean and dashed lines are the 95% credible intervals.

**Fig 6.**
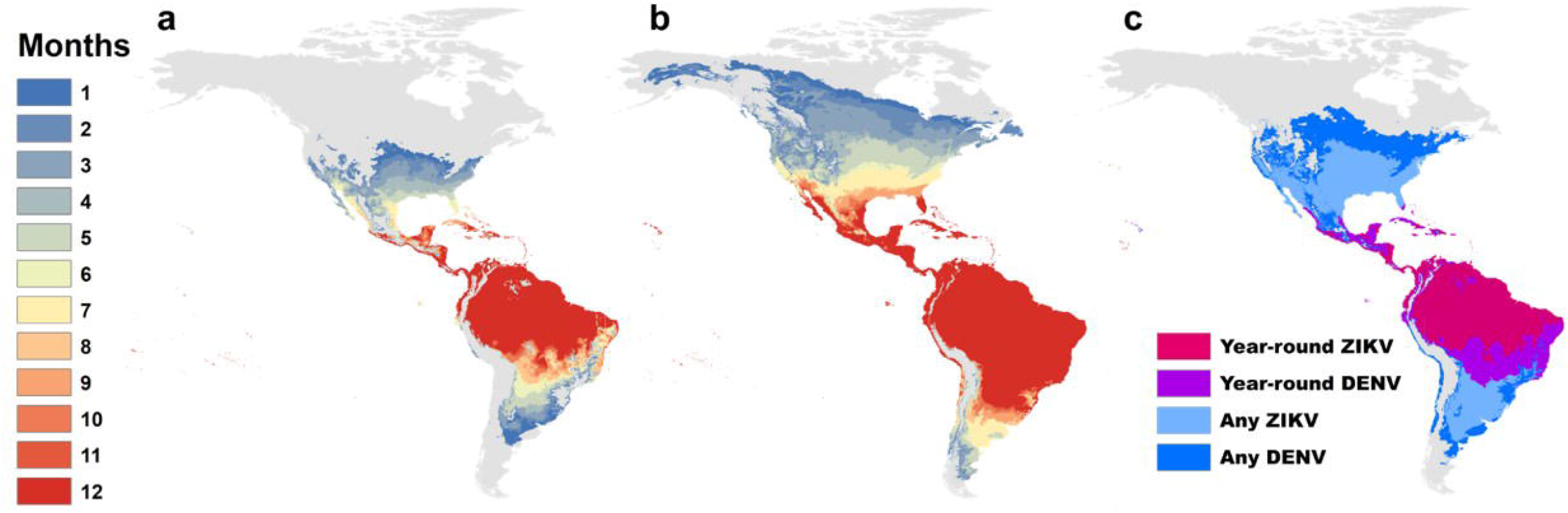
Months of transmission suitability in the Americas. The number of months of transmission suitability (*R_0_*>0) for a. ZIKV derived in this study, and b. Mordecai et al 2017, *Aedes aegypti* only, for median (posterior 50^th^ percentile) models, and c. overlaid all (1-12 months) and year-round (12 months) transmission suitability from a. (light blue, light purple) and b. (dark blue, dark purple), respectively.

While there is some evidence that mosquito longevity varies for virus-exposed versus control mosquitoes, where unexposed mosquitoes had shorter lifespans at near-optimal temperatures (24°C and 28°C; Fig 4 and S3), we did not include this difference in the *R_0_* model for two reasons. First, with limited data to parameterize the low temperature range for survival, we are unable to characterize the differences in the lower end of the thermal response functions in detail. Second, the standard *R_0_* model does not incorporate differences in survival for infected versus uninfected mosquitoes because it assumes that the pathogen is rare and that all mosquitoes are uninfected. For this reason, we fit a single thermal response function for lifespan to the full dataset and used it in the *R_0_* model.

## Discussion

The dynamics and distribution of vector-borne diseases depend on the interplay between the pathogen, the mosquito, and the environment [31]. Temperature is a strong driver of vector-borne disease transmission, and characterizing the thermal range and optimum for transmission is essential for accurately predicting how arbovirus emergence and transmission will be affected by seasonality, geography, climate and land use change. Yet current models of recently emerging arboviruses (e.g. CHIKV [25, 32] and ZIKV [e.g. 6, 7, 9]) are constrained by a lack of data on the thermal sensitivity of key pathogen traits. In this study, we experimentally estimated the relationship between temperature and measures of ZIKV vector competence, extrinsic incubation rate, and mosquito mortality. By incorporating these temperature-trait relationships into an existing mechanistic model, we demonstrate that like malaria [20, 33] and dengue virus [8], ZIKV transmission also has a strong unimodal relationship with temperature.

The effect of temperature on ZIKV transmission is shaped by the complex interaction of individual trait responses of the mosquito and the pathogen with temperature. As studies have demonstrated in other arbovirus systems, temperature significantly affects vector competence [8, 21, 22, 34–39]. We show that temperature has a unimodal relationship with vector competence, with an estimated optimum at 30.6°C and an estimated thermal minimum and maximum of 22.9°C and 38.4°C, respectively (based on posterior median estimates for *T_0_* and *T_m_*). ZIKV infection was limited by different mechanisms at the thermal minimum and maximum. Cool temperatures limited midgut escape and dissemination resulting in a lower proportion of the mosquito population that was infectious. This could be due to temperature effects on mosquito physiology [40], immunity [17, 41–44], and viral binding to specific receptors in the midgut, secondary tissues, and salivary glands [45]. Warmer temperatures, on the other hand, were very permissive for ZIKV infection, resulting in 95% and 100% infection among surviving mosquitoes at 36°C and 38°C, respectively (Fig S2). However, high mosquito mortality resulted in an overall low proportion of the mosquito population becoming infected and infectious (Fig S3). A similar nonlinear effect of cool and warm temperatures on vector competence was observed with *Ae. albopictus* infected with DENV2 [39]. In contrast, Adelman et al. [18] demonstrated that cooler temperatures resulted in increased susceptibility to chikungunya and yellow fever virus due to impairment of the RNAi pathway. However, we only exposed adult mosquitoes to varying mean temperatures, while Adelman et al. [18] looked at the carry-over effects of larval rearing temperature. Both larval and adult temperature variation will likely be important in the field in determining temperature effects on mosquito and pathogen traits comprising arbovirus transmission.

We also observed an asymmetrical unimodal relationship between temperature and the extrinsic incubation rate of ZIKV, with the extrinsic incubation rate optimized at 36.4°C and minimized at 19.7°C and 42.5°C (based on posterior median estimates for *T_0_* and *T_m_*). The effects of temperature on the extrinsic incubation periods of arboviruses and other mosquito pathogens have been extensively studied (dengue virus [39, 46, 47], yellow fever virus [22], West Nile virus [21], chikungunya virus [48], and malaria [49, 50]). Consistent with previous studies, we show that the extrinsic incubation rate of ZIKV increased with warming temperatures, with no infectious mosquitoes observed at 16°C after 21 days post infection and the first infectious mosquito detected at day 3 post infection at 38°C. The extrinsic incubation rate was ultimately constrained at the warmer temperatures due to high mosquito mortality. This is not surprising as metabolic reaction rates tend to increase exponentially to an optimal temperature, then decline rapidly due to protein degradation and other processes [51, 52].

The optimal temperature for mosquito fitness and viral dissemination need not be equivalent, and the impacts of temperature on mosquito mortality relative to the extrinsic incubation rate of arboviruses can have important implications for the total proportion of the mosquito population that is alive and infectious [49, 53]. In our study, mosquito lifespan was optimized at 24.2°C and minimized at 11.7°C and 37.2°C, respectively (based on posterior median estimates for *T_0_* and *T_m_*). The non-linear relationship between metrics of mosquito mortality or lifespan and temperature has also been demonstrated for *Ae. aegypti* [8], *Ae. albopictus* [8, 19] and various *Anopheles* spp. [20, 50]. Despite the fact that the extrinsic incubation period was optimized at a warm temperature (36.4°C), the optimal temperature for overall ZIKV transmission (*R_0_*) was predicted to be cooler (28.9°C) because mosquitoes have a significantly shortened lifespan above 32°C. In contrast, even though mosquitoes are predicted to have relatively longer lifespans at cooler temperatures, the time required for mosquitoes to become infectious (>21 days at 16°C and 18 days at 20°C) may be longer than most mosquitoes experience in the field. As a result, large vector populations may not be sufficient for transmitting the virus if viral replication is inhibited or if the lifespan of the mosquito is shorter than the extrinsic incubation period [54]. One surprising result was that mosquitoes exposed to ZIKV were predicted to live significantly longer at temperatures that optimized mosquito survival as compared to unexposed mosquitoes (37 vs. 87 days at 24°C; 45 vs. 54 days at 28°C). Additionally, the temperature that optimizes mosquito lifespan might also vary between ZIKV exposed mosquitoes (24 °C) and their uninfected counterparts (28°C). If similar trends hold for other arbovirus systems, current modeling efforts may be underestimating virus transmission potential under certain environmental scenarios. If a survival benefit of virus exposure regularly occurs at optimal temperatures across arbovirus systems, estimating mosquito mortality in the field for mosquitoes of different infection statuses and the physiological underpinnings of this response are important areas for future research.

After incorporating the relationships between temperature and vector competence, the extrinsic incubation rate, and mosquito lifespan into a mechanistic model, we demonstrated that ZIKV transmission is optimized at a mean temperature of approximately 29°C, and has a thermal range of 22.7°C to 34.7°C. Because this relationship is nonlinear and unimodal, we can expect as temperatures move toward the thermal optimum due to future climate change or increasing urbanization [55], environmental suitability for ZIKV transmission should increase, potentially resulting in expansion of ZIKV further north and into longer seasons. There is evidence that this is already occurring with warming at high elevations in the Ethiopian and Columbian highlands leading to increased incidence of malaria [15]. In contrast, in areas that are already permissive and near the thermal optimum for ZIKV transmission, future warming and urbanization may lead to decreases in overall environmental suitability [23]. Accurately estimating the optimal temperature for transmission is thus paramount for predicting where climate warming will expand, contract, or shift transmission potential.

By using a mechanistic model originally parameterized for DENV, we also explored a common assumption made by multiple models that DENV transmission has a similar relationship with temperature as ZIKV [6–9]. While the temperature optimum and maximum for *R_0_* changed very little from our previous DENV *R_0_* model, the temperature minimum for transmission increased by nearly five degrees in the updated model (Fig S5). This is mainly due to a higher thermal minimum for both vector competence and the extrinsic incubation rate for ZIKV as compared to DENV (Fig S5 [8]). Differences in the thermal niche of ZIKV relative to DENV or our field derived *Ae. aegypti* relative to those populations assessed in Mordecai et al. [8] could explain this different. There is evidence in a range of invertebrate-pathogen systems (spanning fruit flies, *Daphnia* pea aphids, and mosquitoes) that the effects of environmental variation on disease transmission are often modified by the genetic background of the host and infecting pathogen [38, 56–61]. Thus, more work is required to validate the generalizability of these models.

Our mapped seasonal ranges underscore the impact of a more refined empirical derivation of a pathogen-specific temperature dependent *R_0_*, contrasted with the *Aedes aegypti* dengue prediction of previous studies [6–8]. The higher predicted thermal minimum for ZIKV resulted in a contraction in the areas of the Americas where year-round, endemic transmission suitability (12 months only) are predicted to occur. This area corresponds to a change of approximately 4.3 million km^2^ in land area (Fig 6). Additionally, this higher thermal minimum contributes to a reduction in the overall estimated suitability for ZIKV transmission (all 1-12 months of transmission) resulting in an estimated difference of 6.03 million km^2^. In particular, in the Florida peninsula where the primary focus of ZIKV cases within the U.S. occurred, our updated model (the median model – 50^th^ percentile posterior) now predicts only around six months of temperature suitability during the year (Fig 6) vs. almost year-round as predicted by a previous temperature-dependent Ro model parameterized on the *Ae. aegypti*-DENV system [8]. This contrast in seasonal suitability where ZIKV established in the USA is striking, and emphasizes the value of increasing empirical data and reexamining these types of model, as the capacity to do so becomes possible, in the face of an emerging epidemic.

Finally, although we estimated the effects of mean constant temperatures on ZIKV transmission. Yet mosquitoes and their pathogens live in a variable world where temperatures fluctuate daily and seasonally, and temperature-trait relationships have been shown to differ in fluctuating environments relative to constant temperature environments [23, 62–64]. While characterizing trait responses to mean constant temperatures and incorporating these relationships into models of disease transmission is tractable, more effort is needed in validating computational approaches to infer transmission in a fluctuating environment (i.e. rate summation [8, 65]).

Accurately predicting arbovirus transmission will be influenced by variation in other sources of abiotic (e.g. relative humidity, rainfall), biotic (e.g. availability and quality of oviposition and resting habitats), and socioeconomic factors that influence human exposure to biting mosquitoes. However, this is a fundamental first step for empirically defining and validating current models on the environmental suitability for ZIKV transmission, in which temperature will be a strong driver. Understanding the unimodal effect of temperature on emerging arboviruses, like ZIKV, will contribute to accurate predictions about how future land use and climate change could modify arbovirus emergence and transmission through shifts in mosquito microclimate. *R_0_* models have been used as a tool to guide vector-borne disease interventions, and is a comprehensive metric of pathogen fitness. We anticipate, as with other vector-borne diseases [8, 20, 33], that environmental suitability for ZIKV transmission could expand northwards with future warming, but will be more constrained at low temperatures than DENV. We also predict areas that are already at or near the thermal optimum of 29°C to experience a decrease in environmental suitability for ZIKV transmission [20, 23]. Further, land use change that modifies the microclimates mosquitoes experience could have immediate impacts on ZIKV transmission [55], which might explain the explosive spread of ZIKV in urban centers throughout the Americas.

## Acknowledgments

We thank the University of Texas Medical Branch Arbovirus Reference Collection for providing the virus. We thank Dr. Américo Rodríguez from the Instituto Nacional de Salud Pública for providing mosquito eggs. We gratefully acknowledge the members of the Murdock and Brindley labs for thoughtful comments on the project and manuscript. This study was supported by the National Science Foundation, Grants for Rapid Response Research (NSF-RAPID) 1640780. Erin A. Mordecai and Sadie J. Ryan were supported by NSF DEB-1518681. Erin A. Mordecai was additionally supported by the Stanford University Woods Institute for the Environment Environmental Ventures Program. Sadie J. Ryan was additionally supported by the CDC grant 1U01CK000510-01: Southeastern Regional Center of Excellence in Vector-Borne Diseases: the Gateway Program, to SJR. This publication was supported by the Cooperative Agreement Number above from the Centers for Disease Control and Prevention. Its contents are solely the responsibility of the authors and do not necessarily represent the official views of the Centers for Disease Control and Prevention. Any opinions, findings, and conclusions or recommendations expressed in this material are those of the authors and do not necessarily reflect the views of the National Science Foundation.

